# The Global Avian Invasions Atlas - A database of alien bird distributions worldwide

**DOI:** 10.1101/090035

**Authors:** Ellie E. Dyer, David W. Redding, Tim M. Blackburn

## Abstract

The introduction of species to locations where they do not naturally occur (termed aliens) can have far-reaching and unpredictable environmental and economic consequences. Therefore there is a strong incentive to stem the tide of alien species introduction and spread. In order to identify broad patterns and processes of alien invasions, a spatially referenced, global dataset on the historical introductions and alien distributions of a complete taxonomic group is required.

Here we present the Global Avian Invasions Atlas (GAVIA) – a new spatial and temporal dataset comprising 27,723 distribution records for 971 alien bird species introduced to 230 countries and administrative areas spanning the period 6000BCE – AD2014. GAVIA was initiated to provide a unified database of records on alien bird introductions, incorporating records from all stages of invasion, including introductions that have failed as well as those that have succeeded. GAVIA represents the most comprehensive resource on the global distribution of alien species in any major taxon, allowing the spatial and temporal dynamics of alien bird distributions to be examined.

## Background & Summary

The Parties to the Convention on Biological Diversity (CBD) made a commitment in 2002 to develop an adequate knowledge base to address the problem of invasive alien species, including encouraging research on “the history and ecology of invasion (origin, pathways and time-period)”^1^. Despite this, there continues to be an absence of high-quality, spatially and temporally explicit data available on the distributions of alien species. An evaluation of progress towards the CBD’s 2010 targets^2^ highlighted the need for datasets with broader taxonomic and geographic coverage than those that currently exist^3^. Broad taxonomic coverage is necessary because taxa differ in their likelihood of becoming invasive when introduced, and some will pose greater risks to the new environment or entail greater economic costs to eradicate than others. Broad geographic coverage is needed as currently the majority of data on alien species is skewed towards developed nations^4^, and it is difficult to distinguish whether this imbalance is due to a higher incidence of introductions in these regions or just a greater recording effort. In the absence of broad coverage, any pattern apparent in a dataset is inclined to reflect the pattern in recording effort instead of the true global picture.

In order to address this data gap and begin to identify patterns and processes of alien invasions, a novel, spatially referenced, global data set on the historical introductions and alien distributions of a complete taxonomic group is required. Here we present what is, to our knowledge, the largest and most complete global database on alien introductions and distributions for a significant taxonomic group, birds. Birds provide an excellent focal taxon for studies of invasion biology^5^,^6^. The practise of introducing birds is a global phenomenon, and the wide range of motivations for humans to transport bird species outside of their native ranges has led to a diverse selection of bird species being introduced^5^. In addition, birds are taxonomically well-described, and have had their native distributions mapped at the global scale^7^.

This database on alien bird species distributions derives from both published and unpublished sources, including atlases, country species lists, peer-reviewed articles, websites and via correspondence with in-country experts. The underlying data consist of individual records, each concerning a specific alien bird species introduced to a specific location, and where possible with an associated distribution map. The database forms the core of the GAVIA (<u>G</u>lobal <u>AV</u>ian Invasions <u>At</u>las) project.

Between July 2010 and March 2014, 27,723 alien bird records were collated, representing 971 species, introduced to 230 countries and administrative areas across all eight biogeographical realms, spanning the period 6000 BCE - AD 2014. Raw data comprises taxonomic (species-level), spatial (geographic location, realm, land type) and temporal (dates of introduction and spread) components, as well as details relating to the introduction event (how and why the species was introduced, whether or not it is established).

The number and diversity of the records in GAVIA means that this database should provide a representative portrayal of the global distribution of alien bird species. Indeed, GAVIA doubles the number of bird species known to have been introduced, and also doubles the number known to have established viable populations since Long (1981)^8^, the last attempt at a comprehensive catalogue of alien birds^9^. The coverage of the GAVIA database, both geographically (230 countries), taxonomically (∼10% of all bird species) and temporally (anecdotal records from ∼8,000 years ago, detailed distribution records spanning the last 1,500 years), illustrates the extent of alien bird introductions and spread, and the breadth of available information relating to them. GAVIA represents the first time these data have been collated and compiled into one database, and distribution maps have been created. It is therefore arguably the most comprehensive resource on the global distribution of alien species in any major taxon.

The data contained within GAVIA constitute a large evidence base for the analysis of spatial and temporal patterns in alien bird distributions, and will be an important resource for scientists interested in understanding the invasion process. Multiple publications have already arisen from these data^6,9,10,11,12,13^, however there are still many aspects yet to be explored. Overlaying the GAVIA data with datasets of environmental variables or species attributes provides a wealth of additional analytical possibilities, and should significantly increase the breadth of our understanding of invasions as a result. GAVIA could also help conservation bodies and policy makers to understand where and why invasions are continuing to occur, and so ultimately contribute to efforts to stem the process and ameliorate its impacts.

## Methods

### Data searches

To ensure that equal effort was assigned to gathering data from all regions of the globe, and for all species, the globe was divided into the following regions: North America, Central America and the Caribbean, South America and Antarctica, Europe, Africa, Central Asia, Southeast Asia, Australasia and Oceania. Searches were then conducted for each region in turn, and more general searches were undertaken in order to capture data from global resources.

Online searches of published literature were conducted using Google Scholar, Science Direct, JSTOR and Web of Science. One by one the words ‘invas*’, ‘introduc*’, ‘alien’, ‘exotic’, ‘non-native’, and ‘establish*’ were used to search the literature, together with the name of the region, or the names of the individual countries within that region. Initially these broader invasion biology terms were used in order to pick up more general multi-species studies. Subsequently, the words ‘bird’, ‘avian’ and ‘ornitholog*’ were included in turn. For widely known introduced bird species, a search was conducted using both their binomial and common name(s), e.g. ‘*Acridotheres tristis*’, ‘Indian myna’, ‘common myna’. The reference lists from these articles were searched to identify further papers or books which may have contained useful information.

If the papers or other sources identified from these searches could not be downloaded digitally, then the COPAC national library catalogue (http://copac.ac.uk) was used to identify libraries at which hard copies could be obtained. Hard copies of references came from the Zoological Society of London’s library, the Natural History Museum libraries in London and Tring, Oxford University’s Bodleian and Ornithological (Alexander) libraries, and the British Library. During visits to the libraries listed above, the zoological and ornithological sections were also searched, as well as every country or taxon-specific bird guide, in addition to books relating to invasion biology. As well as articles written in English, articles written in Spanish, German and Mandarin - languages in which one or more of the team of compilers were proficient - were also considered. In addition to published literature searches, the same search terms described above were entered into Google to identify relevant online datasets or country-level species lists which may have contained records of alien bird species.

The names and contact details of people or organisations that were potential sources of information were gleaned from the above literature, and websites (www.europe-aliens.org/expertSearch.do, www.birdlife.org/worldwide/partnership/birdlife-partners) were also used to identify possible experts on alien bird distributions. These contacts were emailed by or on behalf of EED to inform them about the GAVIA project, and to enquire as to whether they knew of any alien bird resources based in their region, or if they knew of anyone conducting similar work. In total, 603 experts from 155 countries were contacted, and useful replies were received from 201 experts from 85 countries. These personal communications proved to be an invaluable resource providing unpublished data and local information, as well as suggestions of obscure published works, or further contacts interested in similar issues.

### Criteria for data inclusion

To be included in the database, records had to meet *both* 1) and 2) from the following criteria, and then *either* 3) or 4) or 5):

1. The record related, at the minimum, to the country level presence of an alien bird species
2. The record identified, at the minimum, the genus to which the bird concerned belongs
3. The record referred to a bird species that had been introduced (either purposefully or accidentally) into an area outside of its native range
4. The record referred to a bird species that had spread to a new area beyond its native range from an adjacent introduced population
5. The record referred to a bird species introduced into an area outside of its historical native range for the purposes of conservation

Records excluded from the database included:

- Single escapees, for example, the blue-and-yellow macaw (*Ara ararauna*) seen flying down Berkhamsted High Street by TMB.
- Migratory bird species occurring as vagrants.
- Records referring to bird species that have naturally expanded their native range into areas immediately adjacent to their original range (e.g. the collared dove (*Streptopelia decaocto*) in Europe).

Including such a broad array of data means that GAVIA contains information on all introduction events, and not only those resulting in establishment. This will enable future research to incorporate a measure of colonisation pressure (*sensu*^14^) into analyses, a variable that is an important determinant of alien species richness (Dyer *et al.* in review) but is usually unavailable.

### Database design

GAVIA was compiled in the programme Microsoft Access 2010. Each entry in GAVIA corresponds to a single record of a single species recorded as introduced and non-native in a specific location as published in a single reference. The data fields of the GAVIA database are described in Table 1. For the sake of minimising repetition, it was decided at the design stage that only ‘new’ data on the actual introduction and invasion events themselves would be collated in GAVIA. Data that would be useful for analytical purposes but which was already recorded elsewhere (e.g. life history data) would not be repeated there. To minimise errors and to reduce the size of the resulting database, supplementary datasets for taxonomy and geographical regions were embedded, and linked to the database via ‘look-up’ tables. This meant that each taxonomic or geographical name was selected through a drop-down list and did not have to be typed repeatedly. This not only significantly reduced the size of the database, and therefore the necessary storage capacity, but also reduced the likelihood of inputting errors. The resulting selection was recorded in the database as an ID number which relates to the species name or country.

**Table 1.** Data fields in GAVIA. ‘Field Name’ shows the GAVIA column headings, ‘Field Type’ denotes what kind of data entry is possible for that field, and ‘Description of Contents’ describes what kind of information is recorded in that field. For Field Type, an ‘Autofill box’ is one which is filled in automatically once a new record is created. For example, each new record is awarded its own unique ID number which cannot be chosen or edited. When a binomial is selected, the respective unique species ID and common name boxes are also automatically filled in and cannot be changed or edited unless a new binomial is selected. A ‘Look-up table’ field type means that the information in that box has been selected from an embedded table, for example the taxonomic list or the GADM country list. In other words there are a finite number of selections from which to choose, and the contents of these cells cannot deviate from the contents of the respective look-up tables. A ‘Free Text’ or ‘Free Number’ box means that the data compiler can freely enter whatever text or number that they wish. A tick box provides the compiler with a certain number of selections, for example island type, and the compiler then ticks the relevant box. An ‘EndNote citation code’ relates to the full references recorded in the GAVIA EndNote reference list.

| Field Name | Field Type | Description of Contents |
| --- | --- | --- |
| RecordID | Autofill box | A unique number for that particular record. Each corresponding individual map also carries this number. This number never changes, even if previous records are deleted. |
| SpeciesID | Autofill box | A unique number for each individual species. |
| Order | Autofill box | The Order to which the species belongs, as per the taxonomy accepted by the IUCN and BirdLife. |
| Family | Autofill box | The Family to which the species belongs, as per the taxonomy accepted by the IUCN and BirdLife. |
| Genus | Autofill box | The genus of that species, as per the taxonomy accepted by the IUCN and BirdLife. |
| Species | Autofill box | The species name of that species, as per the taxonomy accepted by the IUCN and BirdLife. |
| Binomial | Look-up table | The binomial of that species, as per the taxonomy accepted by the IUCN and BirdLife. |
| CommonName | Autofill box | The common name of that species, as per the IUCN and BirdLife. |
| CountryName | Look-up table | The name of the country in which that record occurs as per the GADM designations. |
| AreaName1 | Look-up table | The first sub-level down from country, e.g. region/state, in which that record occurs, as per the GADM designations. |
| AreaName2 | Look-up table | The second sub-level down from country, e.g. sub-region/city, in which that record occurs, as per the GADM designations. |
| LocationDescription | Free text box | A specific description of where the record occurs, if it cannot be selected from AreaName 1 or 2. |
| Realm | Look-up table | The biogeographical realm in which that record occurs, as per the Olson *et al.*<sup>17</sup> delineations (Figure 1). |
| Island | True/False | Whether the record occurs on an island or not. |
| LandType | Look-up table | The type of land that the record occurs on, choices being mainland, continental island or oceanic island. |
| CPrecord | True/False | Whether the record represents a colonisation pressure (CP) location, i.e. a specific location where the species was introduced for the first time, as opposed to a location to which the species has spread. |
| IntroducedDate | Free text box | The date that the species was first introduced (if known), written exactly as found in the reference, e.g. 'late 17th century'. |
| IntroducedDateGrouped | Free number box | The date that the species was first introduced (if known), converted to a number, e.g. 'late 17th century' would become 1690. Guidelines were produced to aid this, so that all transformations were consistent (Table 3). |
| MappingDate | Free number box | The date that the map which corresponds to that particular record represents. For example, the introduced date will stay the same for all individual records from that reference, but as the species spreads over time, the mapped date will change to reflect the newly colonised areas. If there are no dates mentioned at all within the reference, then the date that the reference was published is used as the default mapping date. |
| ReferenceDate | Free number box | Rarely used. If there is no date of introduction recorded, but the reference referred to is a significantly 'old' date, then this is recorded so that it is at least an indication of how long the species has been present in that region. |
| StatusCat | Look-up table | The status of the species in that record, e.g. established, died out etc. (Table 2). |
| IntroMethod | Look-up table | How the species was introduced. For example it was released, or it escaped etc. |
| IntroPurpose | Look-up table | Why the species was introduced. For example it escaped from a zoo, or was released for hunting purposes. |
| TaxonomicNotes | Free text box | Any taxonomic information relevant to that record. |
| Notes | Free text box | Relevant additional notes relating to the record that cannot be entered by using one of the above fields, e.g. it might specify numbers of birds released, or specific paths of species spread etc. |
| RangeMap | Free text box | Whether or not the record has a corresponding distribution map. Either Mapped or Not Mapped. If Not Mapped, it means that it will never be mapped, as the data is deemed too broad scale or vague. |
| Reference | EndNote citation code | Where the information was found, this links to the full list of GAVIA references [Data citation 1]. |
| CompilerInitial | Look-up table | The initials of the person responsible for compiling that record in the database. |

The full bird taxonomy used in GAVIA was that used by the International Union for the Conservation of Nature (IUCN) Red List of Threatened Species (www.iucnredlist.org, downloaded August 2010). The country and regional designations used in GAVIA were downloaded from the Global Administrative Areas (GADM) database (www.gadm.org downloaded August 2010). References were recorded using EndNote citation software (version X4, Thomson Reuters 2010). In a further effort to reduce human error and save computational space, only the first surname, year and EndNote ID code were recorded in GAVIA, which could then be linked back to the full reference in the EndNote database.

Six categories were used to describe the invasive status of each alien species, and definitions of these are provided in Table 2. These categories were chosen to cover all of the ways in which an alien species may be described as being present in a location. An ‘Unknown’ category was necessary as sometimes, even after communicating with experts, it was not possible to assign a species’ status in a certain area to one of the other categories. The opportunity exists to update these cases if and when their status can be clarified.

**Table 2.**
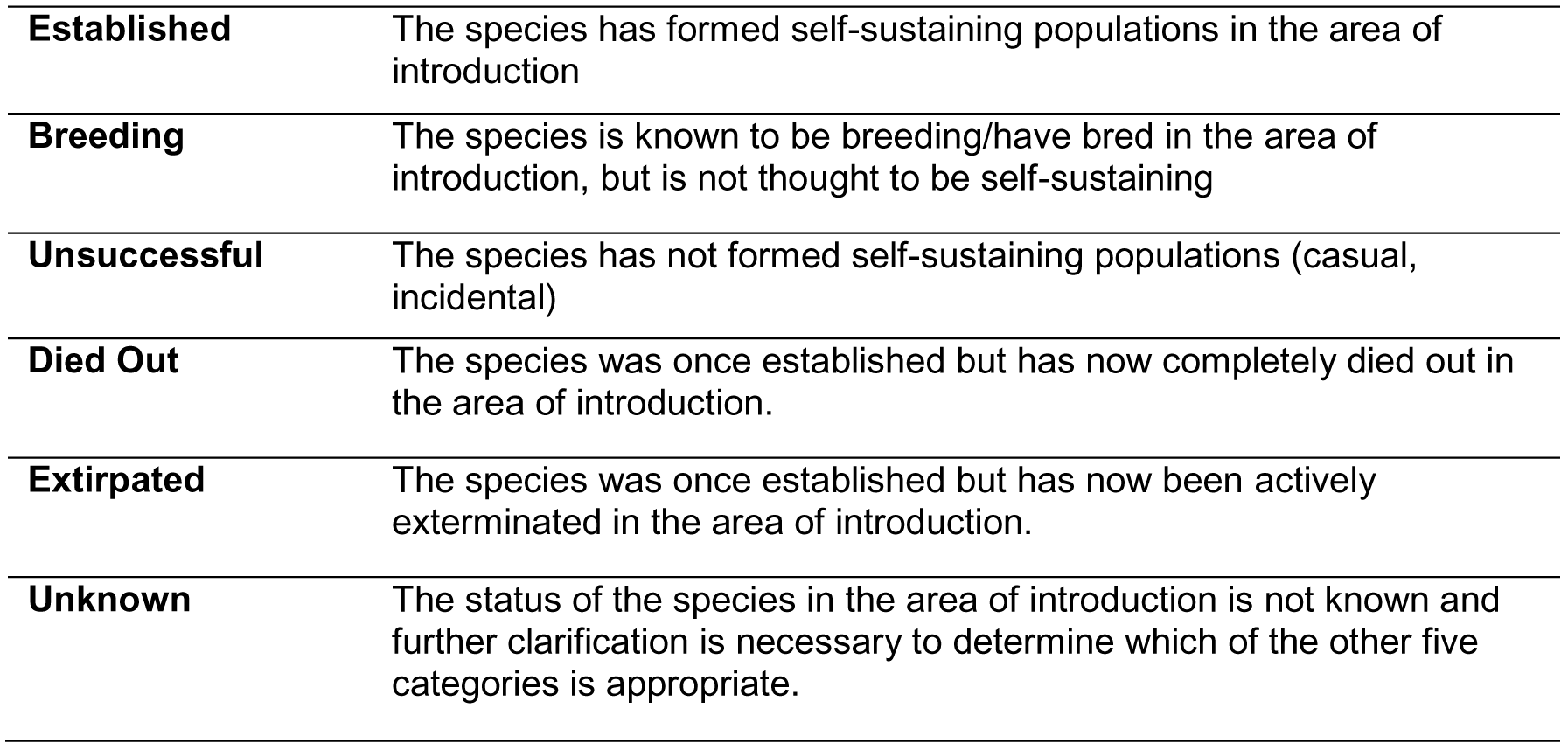
Definitions of alien status categories in GAVIA.

Table 3 demonstrates how dates of introduction were recorded in the GAVIA database. Often in the literature, a date is approximated, or described in a way that is not a four-digit year. In order to maintain the integrity of the reference, the date was first recorded exactly as given in the reference (e.g. ‘early 1700s’). To make the date usable in later analyses, it was also converted to a four-digit number (in the preceding example, this would be 1710) (Table 3). All converted dates were Anno Domini, although four records had dates of introduction earlier than 1000AD, and were consequently converted to three digit numbers. All records with dates of introduction Before the Common Era (BCE) were too vague to convert to a usable date. These guidelines ensured that all data compilers recorded dates in the same fashion.

**Table 3.** Guideliness used for converting the introduced date given in the reference into a whole number.

| Date given | Grouped<br>(converted) date | Rule |
| --- | --- | --- |
| 1912 | 1912 | Use the four digit number as given |
| c.1890 | 1890 | Use the four digit number given |
| 1777-1778 | 1777 | Use the earliest date in the range |
| 1930-1940 | 1935 | Use the midpoint of the range |
| C18th | 1750 | Use the midpoint of the century |
| early C18th | 1710 | Use the date 10 years into the century |
| mid C18th | 1750 | Use the midpoint of the century |
| late C18th | 1790 | Use the date 10 years before the end of the century |
| 1800s | 1850 | Use the midpoint of the century |
| c.1800s | 1850 | Use the midpoint of the century |
| 1990s | 1995 | Use the midpoint of the decade |
| early 1700s | 1710 | Use the date 10 years into the century |
| mid 1700s | 1750 | Use the midpoint of the century |
| late 1700s | 1790 | Use the date 10 years before the end of the century |
| early 1990s | 1991 | Use the first year of the decade |
| mid 1990s | 1995 | Use the midpoint of the decade |
| late 1990s | 1999 | Use the last year before the end of the decade |
| 1980s-1990s | 1990 | Use the midpoint of the two decades |
| <1965 | 1964 | Use the date immediately before the date given |
| >1970 | 1971 | Use the date immediately after the date given |

### Data entry

In total, seven data recorders were involved with entering data into the GAVIA database including EED, four interns and one project assistant, plus a technician who worked for TMB in 2006/7. To maximise uniformity in data entry, all data recorders were given thorough and consistent training, and each was provided with a set of database guidelines. In addition, spot checks were regularly carried out on all database entries, and weekly meetings of the GAVIA team were held to address inconsistencies.

At the time of data collection and entry, all information was entered into the database exactly as it was described in each reference, with as much information extracted as possible. Multiple records from different authors who had recorded the same information were still included in the interests of completeness.

An Access Database form was created to standardise data entry, and this also enabled multiple members of the team to enter data simultaneously. This form was divided into three sections: Taxonomy, Distribution and Introduction. Where available, the following data were entered into the GAVIA database for each record under each section tab (see Table 1 for full details):

#### Taxonomy tab

1. The species’binomial was selected fromadrop downlist, and this then automatically filled in the appropriate Order, Family, Genus, Species, species ID, common (English) name, and any synonyms.
2. A free text box titled ‘Taxonomic Notes’ allowed the complier to enter any additional information regarding the taxonomy of the species in question, for example if it was thought to be a certain subspecies, or if the identification was uncertain.

#### Distribution tab

3. The drop down boxes ‘Country’, ‘Area Name 1’, and ‘Area Name 2’ are the country, state and sub-state level delineations available for selection by the compiler. These areas match up to the GADM spatial layers used in the distribution maps relating to each database record.
4. The free text box ‘Location Description’ was used for additional information regarding the location of the record. For example, it may specify a location not included on the GADM list, or it could provide additional directions such as ‘the area of National Park between town A and town B’.
5. The compiler then selected the biogeographical realm within which the record lay, and also recorded the land type (mainland, oceanic island or continental island), and selected the ‘Island’ tick box if the record occurred on an island of either type.
6. The ‘RangeMap’ box was used to identify whether or not that record contained enough detail to be converted into a distribution range map. At the data entry stage, this box was also used to record whether or not the reference included a distribution map of the species, in which case it was photocopied or printed and stored for later use.

#### Introduction tab

7. The introduction status of the species was selected from a drop down list (Table 2).
8. There were four different date boxes available to the compiler. The ‘Introduced Date’ is the date exactly as recorded in the reference. ‘Grouped Date’ is the introduced date converted to a whole number (if necessary) using the standardised system as described in Table 3. ‘Reference date’ is rarely used, but useful if the record does not include a date of introduction, yet the reference in question is sufficiently old enough to warrant the inclusion of the publication date as an indication of timescale. For example, if the reference was written in 1910, even if it does not state a specified date of introduction it is possible to deduce that the bird was present in that location over a hundred years ago. ‘Mapping date’ refers to the date of any associated distribution map(s). For example, a source may describe a species as having been introduced to a location in the year 1900, but also records that the species had spread to a much larger range size by the year 1950. In this case, two records would be created, resulting in two distribution maps. The first record would have both the date of introduction and the mapping date as 1900, and the map would relate to the distribution of the species at this time (i.e. the location of introduction). The second record would also have the date of introduction as 1900, but the mapping date would be 1950, and the associated map would relate to the subsequent (larger) distribution. If there were no dates mentioned at all within the reference, then the date that the reference was published was used as the default mapping date.
9. The free text ‘Notes’ box was for recording additional relevant information, for example details of spread, or an estimate of population health.
10. Under ‘Method of Introduction’ and ‘Reason for Introduction’, tick boxes allowed the compiler to record how and why the species was recorded, if this information was available.

The Access form acted as an entry portal for data, but the resulting records were stored in an Access table, with each selection from the drop down menus stored as a number; the only text stored is from the free text boxes. This reduced the size and complexity of the database, and reduced the likelihood of errors. An Access query can be run to extract specific information, or to view the entire database in its readable text format.

Where a reference provided information for multiple species or countries, individual records were created for each species-country pair. Information sent to us in email form from experts was recorded in the Endnote library as ‘pers. comm.’ and entered accordingly into the main database.

### Taxonomic names and classification

It was necessary to be able to identify taxa in the database as accurately as possible, and without losing any information. It was also necessary to be able to place each species within the avian phylogeny. Therefore, we required a stable and authoritative resource for nomenclature, which included species whose status may be unclear. The taxonomy used in GAVIA was thus based on that agreed by the IUCN Red List of Threatened Species at the time of database creation (2010).

Two species with records in GAVIA were not included on the IUCN taxonomy: the Javan myna (*Acridotheres javanicus*) and the Barbary dove (*Streptopelia risoria*). The taxonomy of these species is in dispute^15,16^, but as there were substantial records of individuals assigned to these taxa being introduced, the decision was made to add their names to our taxonomic list. If in the future their species status is agreed upon then the records can be updated accordingly.

Where a species name stated in a reference was a synonym for one included in the IUCN taxonomy, the accepted species name was selected on the Access form, and the synonym used in the reference was written in the ‘Taxonomic Notes’ section. Where a subspecies was mentioned in the reference, the record was listed under the species name, and the subspecies was also recorded in the ‘Taxonomic Notes’ section. There are 11 records in the GAVIA database with no attributed species name; these records are excluded.

The use of a drop-down list for selecting the species name on the data entry form, with the higher taxonomy then automatically entered, resulted in minimal errors and inconsistencies when inputting species names. Any typographical errors in the original reference (e.g. misspelling of species names) were again recorded in the ‘Taxonomic Notes’ text box.

### Biogeographical coverage

Alien bird records were compiled for 230 countries and administrative areas from all biogeographical realms (although only offshore islands from the Antarctic realm have records - there are no records (yet) for the Antarctic continent). Realm delineations followed those set out in Olson *et al.*^17^ (Figure 1). A concerted effort was made to identify any alien birds introduced to regions where data was deficient (Figure 2), so we are confident that we can rule out a lack of effort as the reason for the lack of records. However, it is not known whether it is actually the case that no alien birds have been introduced to these places, or whether they have but either no one has recorded them, or these records have not yet found their way into the public domain.

**Figure 1.**
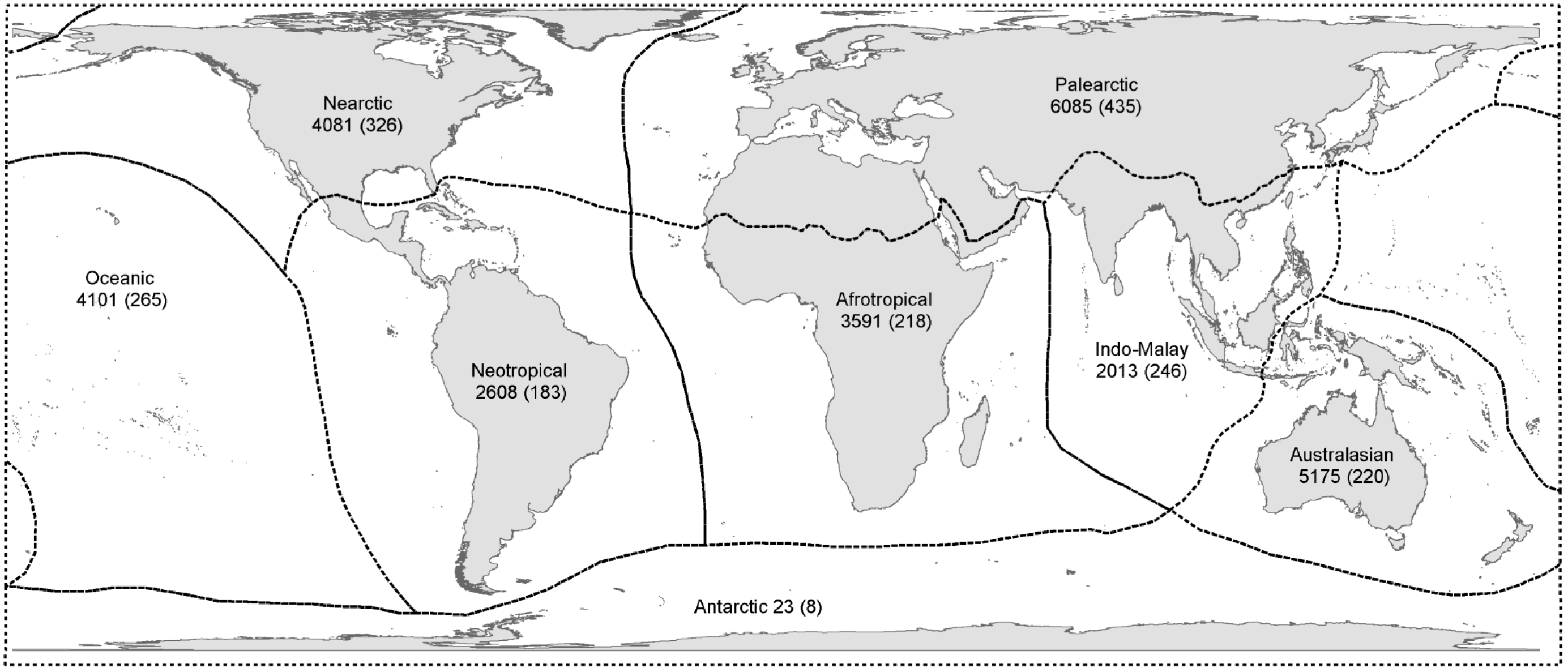
The 8 biogeographical realms used in Olson *et al.*^17^, and which were followed by GAVIA for the purposes of assigning alien ranges to realms. The first number is the number of records in GAVIA for each realm, and the number in brackets is the number of species recorded as being introduced in each realm.

**Figure 2.**
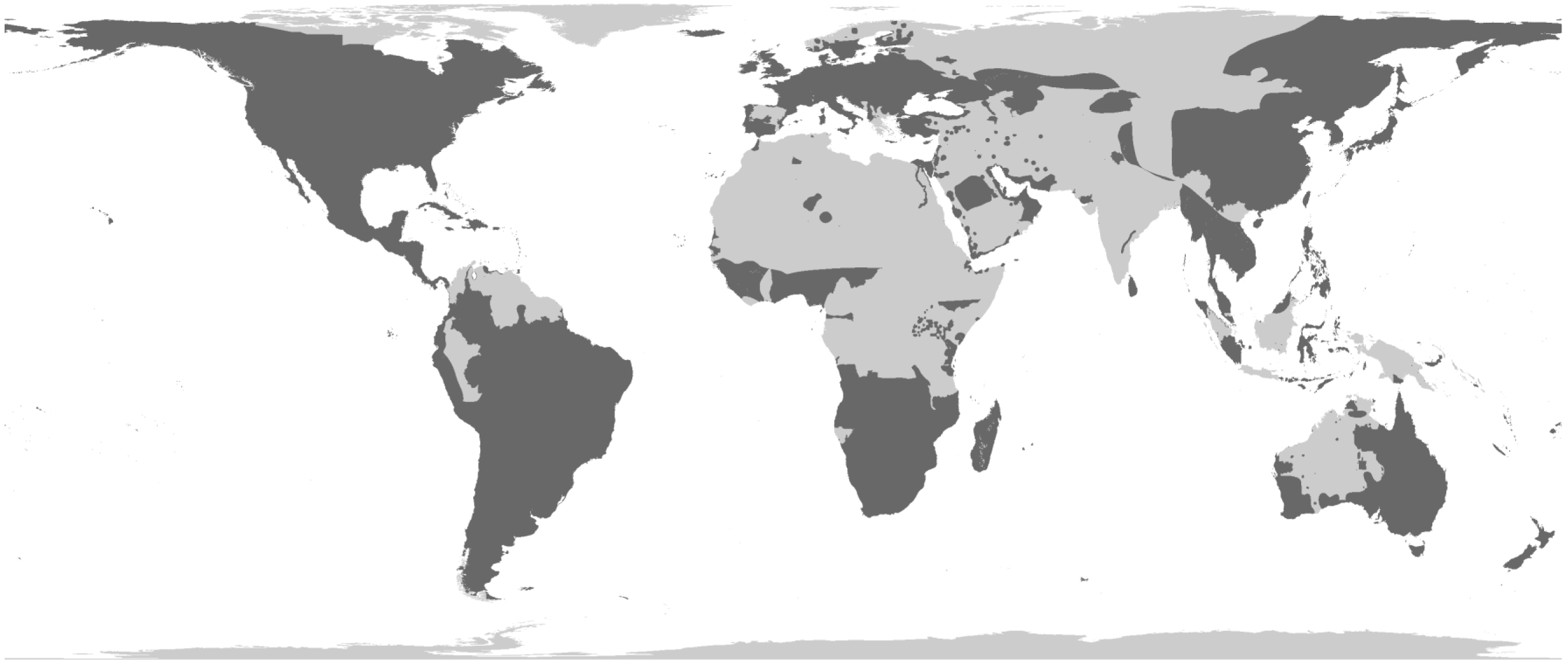
The global distribution of those records in GAVIA that contain sufficient information to have been converted into distribution maps. These include all status categories, so introductions that have both succeeded and failed.

In order to maintain continuity, the list of country units defined in the GADM database was used in the GAVIA database (‘Country’, ‘Area Name 1’ and ‘Area Name 2’), and the corresponding GADM GIS layers (downloaded from www.gadm.org) were used to produce the resulting range maps.

The GADM GIS layers are at a very fine scale, with extremely detailed borders, coasts and island groups. This inevitably led to a considerable increase in the computational memory and storage space required for the maps, and more importantly the processing time for analysis. However, this level of detail was deemed necessary as many alien bird species have been introduced to islands or coastal areas, locations which are simply missing from lower resolution GIS layers. Had a coarser scale base map been used, not only would it have proved difficult to map some of these coastal or island records, but any subsequent analysis involving range size calculations would have been inaccurate.

### Distribution range maps

Introduction records were converted into distribution maps using the software ESRI ArcGIS version 9.3^18^. All records containing a high enough level of detail to create an accurate estimation of distribution range were converted into maps, regardless of alien status. All team members involved in this activity received 2-3 days of training beforehand using the training manual created for internal use at the Zoological Society of London^19^.

In addition to this training, team members received a set of guidelines to follow, and a random sample of distribution maps created by the team each week would be spot-checked to identify any errors or inconsistencies. Any problems were worked through at weekly meetings. This was to ensure, as far as possible, that all team members created distribution maps in a uniform manner.

One of the anticipated problems with having multiple team members accessing the GAVIA database at the same time was the risk of them simultaneously editing the same record, such that one entry would overwrite the other. To prevent this from happening, each member of the team was assigned their own Access query which they could use to extract data from the database. A normal Access query enables the user to view a subset of information from a database, but the data cannot be edited through the query. If a team member did want to open or edit the main Distribution table containing the raw data, they first had to check verbally that no one else was using it or had it open on their screen. In order to keep the team’s files and folders as consistent and logical as possible, all team members followed the guidelines provided to them and adhered to a strict system of file and folder labelling and backing-up.

The website www.geonames.org was used to identify latitude and longitude points for place names, so that they could be plotted. If a hard copy map existed then it was scanned and georeferenced. If the location description only provided information for a single city or point then a 10km buffer was created around it in order to produce a range polygon. Each map file was labelled with the species’ name and record ID. Once all records for a species were converted into range maps, the files were merged together and combined with the previously created attribute table (containing all of the data for that species extracted from the GAVIA database) and saved as a single shapefile uploaded into the main GAVIA folder.

Some records in GAVIA needed to be split before they could be mapped. For example, a record may have stated how the distribution of the species had changed over time. In such cases, multiple maps needed to be created to plot this change. Conversely, some records in GAVIA were deemed not to contain enough detail to warrant conversion into distribution maps. It was important that the resulting distribution maps were as detailed as possible, but were also mapped to a comparable level of detail. If the record only stated the country in which the species was introduced, without further specification of location, then it was recorded as being ‘Not mapped’ in the RangeMap box. Exceptions to this rule were if the country was particularly small (e.g. Singapore, Hong Kong), or if it was a small island (e.g. the majority of the Pacific islands).

Distribution maps were created to the minimum possible range size so as to not overestimate a species’ distribution. When combined, the distribution maps represent the species’ Extent of Occurrence rather than Area of Occupancy^20^, and the species are unlikely to be extant in every part of their mapped range, particularly those areas where their status is not established. The distribution maps were projected using the World Behrmann equal area projection so that accurate range size estimates could be calculated.

### Data summary

Once the records were converted to distribution maps, areas with relatively low numbers of alien birds or no recorded introductions included areas close to the poles (Greenland, northern Russia, far northern Europe, northern Canada, Antarctica), deserts (parts of the Sahara, western and central Australia, the Gobi desert, the Arabian desert), mountainous areas (parts of the Andes and the Himalayas), and parts of the tropics (northern South America, central Africa, and parts of Indonesia, Borneo and Papua New Guinea) (Figure 2). For those records where a land type was assigned, 44% related to oceanic islands (12,203 records), 40% related to mainland locations (11,133 records) and 16% to continental islands (4,263). The best-represented biogeographical realms are the Palearctic (6,085 records, 22% of all records, 435 species), Australasian (5,175, 19%, 220 species), Nearctic (4,081, 15%, 326 species) and Oceanic realms (4,101, 15%, 265 species) (Figure 1). Four countries have more than one thousand records each: the United States (6,158), New Zealand (2,464), Australia (2,363), and the United Kingdom (1,631).

There are records in GAVIA of birds being transported to areas outside of their native distributions *c*. 8,000 years ago (Red Jungle Fowl (*Gallus gallus*)^21^), such that the earliest record is from ∼6000 BCE. However, the earliest record for which there is enough detail for a distribution map to be created is from 500 AD. The most recent date of first introduction (as opposed to the ‘Mapping Date’ or date of spread) is 2010. Therefore, the records in GAVIA with a first date of introduction at a resolution suitable for mapping span 1,510 years.

The cumulative number of records in GAVIA increases steadily until 1850, at which point there is a step-change and the cumulative number of records increases by an order of magnitude over the following 150 years (Figure 3a). An almost identical pattern is apparent in the cumulative number of alien bird species recorded in GAVIA, although on a different scale (Figure 3b). When plotted together, it is possible to see that the number of records and the number of species do indeed increase in parallel, demonstrating that in the last 150 years in particular, more people have been recording a greater variety of alien bird species (Figure 3c). The number of records in GAVIA for each year also demonstrates an increase in recording effort over time.

**Figure 3.**
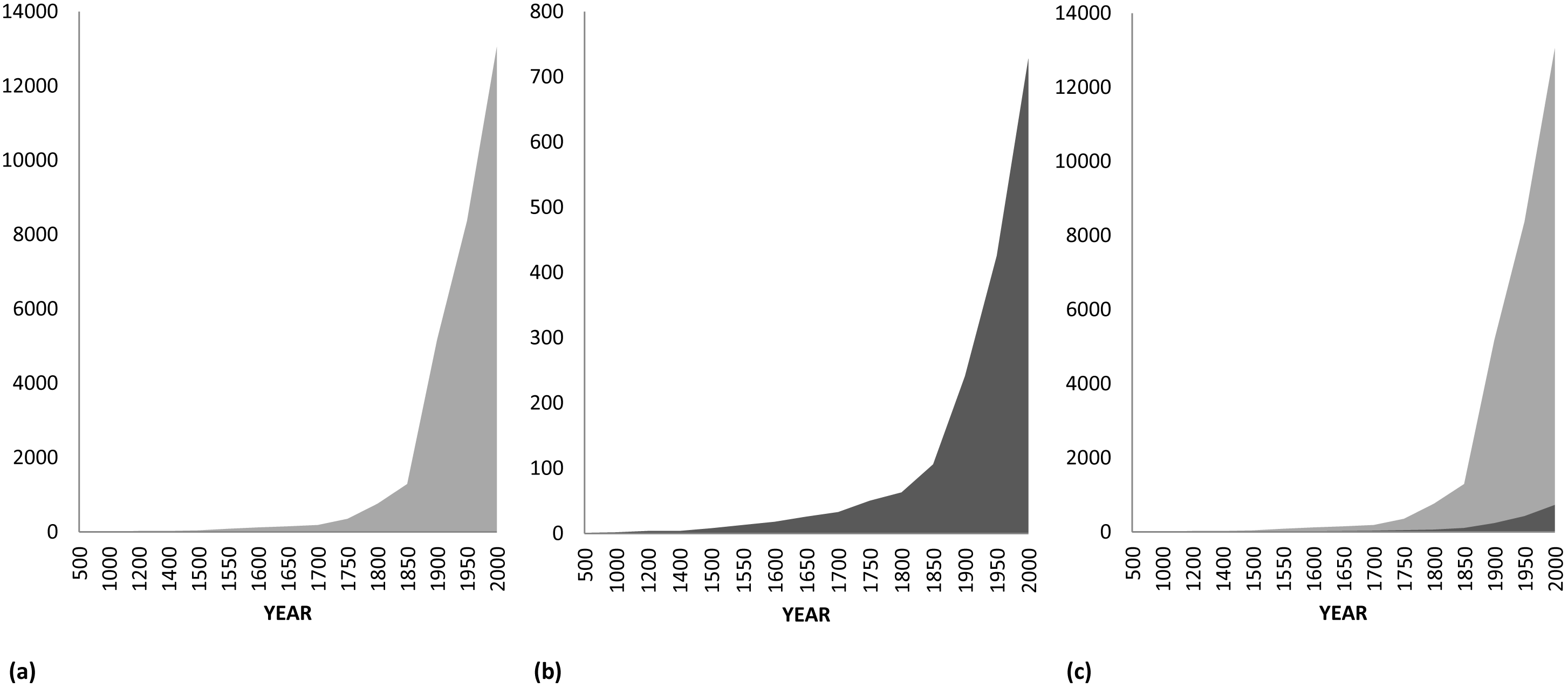
**(a)** The cumulative increase in the number of records in the GAVIA database over time; **(b)** the cumulative number of species recorded in the GAVIA database over time; and **(c)** both the number of records and species plotted together. Light grey is the number of records, dark grey is the number of species.

The bird families with the highest numbers of species records are the parrots (Psittacidae: 131 species recorded), and ducks, geese and swans (Anatidae: 92). Seven species have more than five hundred records each in the database: house sparrow (*Passer domesticus*, 1,292 records), common myna (*Acridotheres tristis*, 1,214), rock pigeon (*Columba livia*, 823), rose-ringed parakeet (*Psittacula krameri*, 778), common pheasant (*Phasianus colchicus*, 681), common starling (*Sturnus vulgaris*, 673) and Java sparrow (*Padda oryzivora*, 540). The highest proportion of records in GAVIA relate to established species (13,144 records, 47% of all records), followed by records with an unknown status (9,141, 33%) (Figure 4).

**Figure 4.**
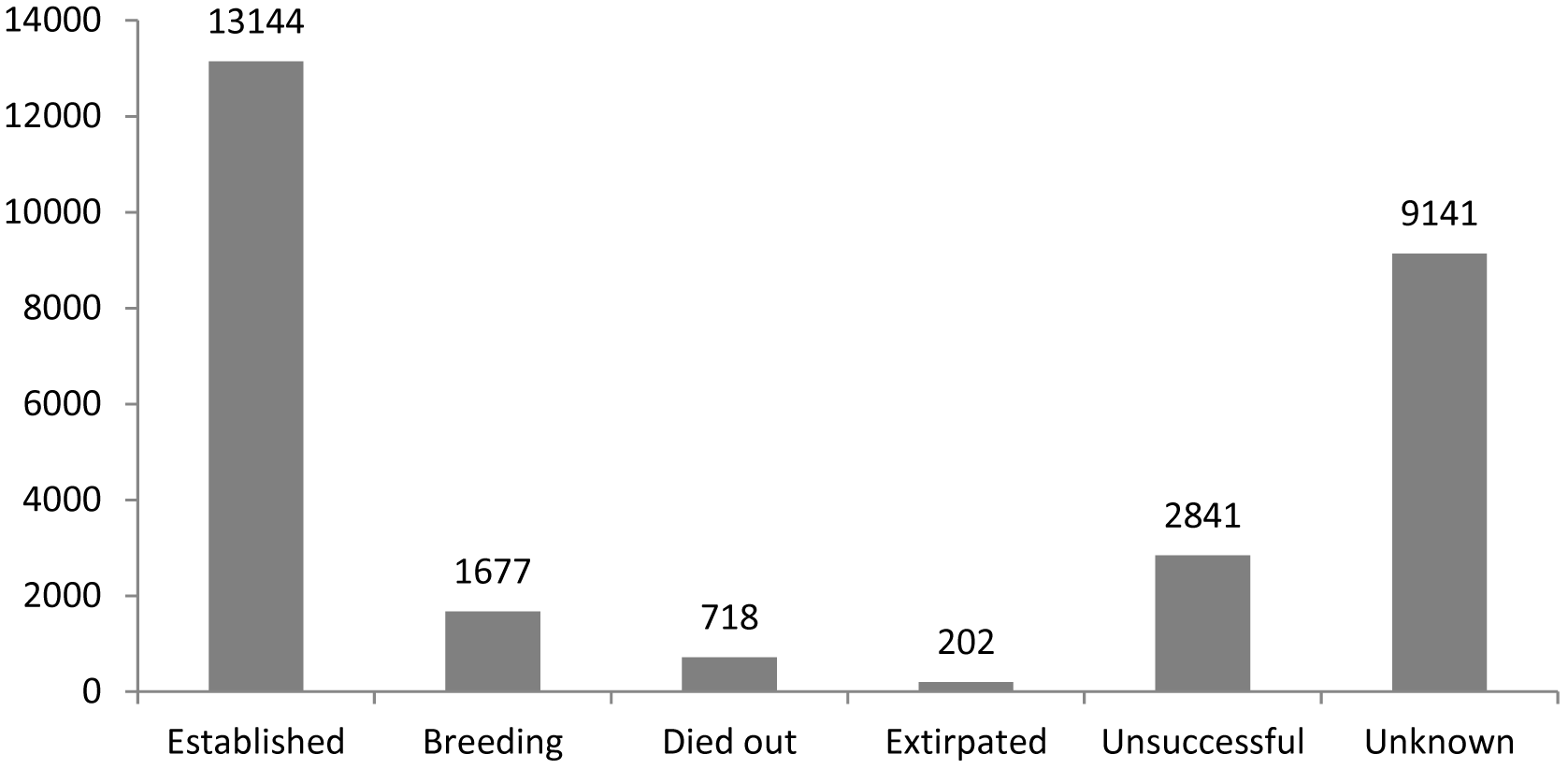
The number of records in GAVIA assigned to each introduction status.

Note that these numbers cannot be used to calculate establishment probability, as established populations are more likely to generate multiple records in the database. The majority of the 971 species in GAVIA have more than one recorded occurrence, for which the outcomes may be different. Thus, 419 species (43%) have an established population somewhere in the world, 464 (48%) have an unsuccessful population somewhere, 236 (24%) have a breeding population, 178 (18%) have a population that was once established but has now died out, and 76 (8%) had a population that has now been extirpated. The status of one or more of the populations of 816 species (84%) is unknown.

### Data Records

All of the following GAVIA data are stored in a *Figshare* data repository [Data citation 1].

The main GAVIA data table is contained in a single comma-separated file (.csv format), entitled ‘GAVIA main data table’. Each row below the header represents a specific alien bird species introduced to a specific location (*n* = 27,723), and the columns (*n* = 28) contain taxonomic, spatial and temporal data, and information on the introduction event.

Full descriptions of the column titles are contained in Table 1, and also in a single comma-separated file (.csv format), entitled ‘GAVIA column names’.

A list of the abbreviations used in the GAVIA data table is contained in the single comma-separated file (.csv format), entitled ‘GAVIA abbreviations’.

The full list of the references referred to in the GAVIA data table is file is contained in the single comma-separated file (.csv format), entitled ‘GAVIA references’.

The species’ range maps are contained in a compressed folder (.zip format), entitled ‘GAVIA rangemaps’, and within this are stored as one ESRI shapefile per species (n = 362) representing the species’ most recently recorded established alien range. Within these shapefiles are attribute tables which contain a unique species ID number and binomial which match up to the species ID number and binomial in the ‘GAVIA main data table’.

### Technical Validation

The final stage of the project required all of the distribution maps to be cross validated against the database. This was carried out by a single team member in an effort to lessen any inconsistencies that might be introduced into the database by different team members. Each species was addressed in turn. Consistency checks were carried out on the records in GAVIA, and then the distribution maps were verified to ensure that they corresponded to the information in the database. In addition to these checks, each species’ alien distribution map was checked against its native range map (representing native global breeding range) extracted from the database used by Orme *et al.^7^.* This was to ensure that there was no overlap, for example regions where a species was native but it had been recorded as introduced or vice versa. Necessary changes were made to both the database and the distribution maps.

### Usage Notes

A common problem with macroecological and invasive species studies is the bias in locations where biologists conduct their research, both geographically and also in terms of habitats which are inaccessible or difficult to survey. This geographical bias is particularly prevalent in single-species studies^22^. Although Europe, the United States and Australia are over-represented in terms of research locales^4^, it is difficult to disentangle whether this is due to a higher number of invasion biologists focussing their studies there, or if it is a justified skew as a result of these areas holding a relatively larger number of alien species. In addition, Pyšek *et al.^22^* found that invasion research seemed to focus on those species that are perceived to have the potential to produce the most economic or ecological harm. Although GAVIA is based on a systematic and thorough search of all the data available from all regions of the world (where possible), there is still the potential for biases due to the intrinsic biases in the available literature. It is likely that there are regions of the world where invasions are continuing to occur without written records being made, and therefore even if the most thorough search of the literature is made, records will still be missed. This potential bias needs to be taken into consideration when conclusions are being drawn from the results presented here.

The use of the GADM layers as a basis for the range maps may have resulted in a small degree of spatial extrapolation of introduction records. For example, if a record states that a species is present in the Australian city of Sydney then the resulting distribution map will encompass the whole of Sydney as delineated by the GADM level 3 layer, although in reality it may only occur in a certain area of the city. This was addressed by producing distribution maps which represented the minimum convex polygon of the range that was described in the record, in order to avoid any unnecessary extrapolation. Where the record was too vague in its spatial description, a distribution map was not created. However, it is possible that for some species, their alien range size may be over-estimated due to this potential extrapolation; as these maps represent Extent of Occurrence^20^, the species is anyway unlikely to be extant in every part of its total recorded alien range, as is also the case with most commonly used native species range maps.

The distribution range map for the common pheasant, *Phasianus colchicus*, is very large and has a substantial detailed border. Therefore for ease of manipulation it consists of two polygons within one shapefile. These polygons do not overlap and together represent the established alien range of this species. All other species distribution range maps consist of one polygon within one shapefile [Data citation 1].

The feral or rock pigeon, *Columba livia*, has a long history of human-mediated global transportation, and as such there is some uncertainty over what constitutes its true native range versus historical introductions. In the GAVIA database, all records where *C. livia* has been referred to as an alien have been included for completeness. However, those records which concern regions where there is some debate over whether *C. livia* is truly alien or not have the caveat: *\*Although described in the reference as alien, part or all of this range may overlap with the species’ native range\** included in the ‘Notes’ column. Where an alien distribution map for *C. livia* has been created, those regions that overlap with the native range used by Orme *et al.*^7^ have been removed so as to prevent the species from being counted as both alien and native in the same location.

## Acknowledgements

The authors thank the 603 in-country experts contacted during the data collation for this database; also Mark Parnell, Victoria Franks, Fiona Spooner, Rebecca Herdson, Elizabeth Jones and Frances Davis for assisting with data collection and map creation. Initial funding for this study was provided by a grant from the Leverhulme Trust (RF/2/RFG/2010/0016) (EED), with additional support from a UCL IMPACT studentship (10989) (EED), and from a King Saud University Distinguished Scientist Research Fellowship (TMB, DWR, EED).

## Author contributions

EED designed the database, conducted data collection, data entry, map creation, curated the data file and wrote the manuscript. DWR assisted with data curation and edited the manuscript. TMB conceived the research, oversaw collation of the database and edited the manuscript.

